# Filter bank common spatial pattern and envelope-based features in multimodal EEG-fTCD brain-computer interfaces

**DOI:** 10.1101/2024.09.15.613144

**Authors:** Alaa-Allah Essam, Ammar Ibrahim, Ashar Zanqour, Mariam El-Saqa, Sohila Mohamed, Ayman Anwar, Ayman Eldeib, Murat Akcakaya, Aya Khalaf

**Author notes:** **Correspondence:** Aya Khalaf, PhD. Equally contributing co-first authors (Alaa-Allah Essam, Ammar Ibrahim, Ashar Zanqour, Mariam El-Saqa, Sohila Mohamed).

## Abstract

Brain-computer interfaces (BCIs) exploit brain activity to bypass neuromuscular control with the aim of providing alternative means of communication with the surrounding environment. Such systems can significantly improve the quality of life for patients suffering from severe motor or speech impairment. Multimodal BCIs have been introduced recently to enhance the performance of BCIs utilizing single modality. In this paper, we aim to improve the performance of multimodal BCIs combining Electroencephalography (EEG) and functional transcranial Doppler ultrasound (fTCD). The BCIs included in the study utilized two different paradigms to infer user intent including motor imagery (MI) and flickering mental rotation (MR)/word generation (WG) paradigms. Filter Bank Common Spatial Pattern (FBCSP) algorithm was used to extract features from the EEG data. Several time series features were extracted from the envelope of the fTCD signals. Wilcoxon rank sum test and linear kernel Support vector machines (SVM) were used for feature selection and classification respectively. Additionally, a probabilistic Bayesian fusion approach was used to fuse the information from EEG and fTCD modalities. Average accuracies of 94.53%, 94.9% and 96.29% were achieved for right arm MI versus baseline, left arm MI versus baseline, and right arm MI versus left arm MI respectively. Whereas average accuracies of 95.27%, 85.93% and 96.97% were achieved for MR versus baseline, WG versus baseline, and MR versus WG respectively. Our results show that EEG- fTCD BCIs with the proposed analysis techniques outperformed the multimodal EEG-fNRIS BCIs in comparison.

## 1 Introduction

Brain computer interface (BCI) is a technology aiming at providing a direct communication channel between the central nervous system and external devices (1,2). Therefore, BCIs can assist individuals suffering from disorders that limits their ability of interaction with the surrounding environment such as stroke, amyotrophic lateral sclerosis, cerebral palsy or spinal cord injury by providing alternative means of communication (3). Other BCI applications include controlling robots (4), prosthetic limbs (5), rehabilitation (6), virtual reality (7) and neurogaming (8).

Both invasive and non-invasive neuroimaging modalities have been used to develop BCIs. Non-invasive BCIs are safer, offering high accessibility, cost effectiveness, and scalability (9,10). Different noninvasive neuroimaging modalities capturing the electrical and metabolic activity of the brain have been used in BCI design including electroencephalography (EEG) (11,12), functional near-infrared spectroscopy (fNIRS) (13), functional magnetic resonance imaging (fMRI)(14) and magnetoencephalography (MEG) (15). Among these modalities, EEG is the most commonly used modality due to its portability, cost-effectiveness and high temporal resolution (16). However, it has a low signal to noise ratio and low spatial resolution. In addition, it is prone to non-stationarities resulting from external electrical interference and internal brain background activity (17). These drawbacks limit the performance of EEG-based BCIs outside laboratory-controlled environments and lead to misidentification of user intent and false-command generation (18).

In order to overcome these limitations, multimodal BCIs employing EEG in addition to other modalities measuring different brain activities such as fNIRS and fMRI were proposed (19). However, EEG-fMRI BCIs cannot be implemented in practice due to their non-portability, high cost, and the need for a highly controlled environment for efficient performance (20,21). fNIRS is the most commonly used second modality in multimodal BCI systems due to its portability and immunity against electrical noise, however, it suffers from low temporal resolution (22–24). Another limitation is that infrared signals can be blocked by the user’s hair (25,26).

Functional transcranial Doppler ultrasound (fTCD) has been suggested as a fast and cost-effective alternative for fNIRS in BCI design (27). fTCD assesses the cerebral blood flow velocity (CBFV) using two ultrasound transducers placed above the zygomatic arch on both left-side and right-side transtemporal windows (28). Recently, we proposed a multimodal BCI combining EEG and fTCD modalities as a faster and more efficient alternative to EEG-fNIRS BCIs (29–33). This BCI employed two different paradigms to induce simultaneous changes in EEG and fTCD through presenting visual stimuli that instruct participants to perform motor imagery tasks (MI paradigm) (30,31) as well as flickering mental rotation (MR) and word generation (WG) tasks (flickering MR/WG paradigm) (29,32).

In this paper, we aim at improving the performance of MI and MR/WG EEG-fTCD BCIs through applying analysis approaches that have not been previously used to analyze multimodal EEG-fTCD data. Moreover, we investigate the contribution of each modality in user intent inference within each paradigm. Three binary classification problems were investigated for each paradigm, including left MI versus baseline, right MI versus baseline, and left MI versus right MI for the MI paradigm as well as MR versus baseline, WG versus baseline, and WG versus MR for the flickering MR/WG paradigm. Common Spatial Pattern (CSP) algorithm is the most effective algorithm for EEG feature extraction due to its computational simplicity and ability to obtain highly separable features in terms of spatial patterns among different motor imagery tasks (34). We employed an extended version of the classical CSP approach named filter bank common spatial pattern (FBCSP) algorithm to extract features from EEG data of both paradigms (35,36). Most studies performing fTCD signals analysis extract features from the raw fTCD data (27,37). In this study, we extracted statistical, temporal, and spectral features from the envelope of the raw fTCD signal which represents maximal cerebral blood flow velocity at each time point. fTCD significant features were selected using Wilcoxon rank sum test. Linear kernel support vector machines (SVM) was used for classification. Instead of concatenating the extracted features vectors from both EEG and fTCD modalities, a probabilistic Bayesian fusion approach was implemented to generate a joint decision from EEG and fTCD evidences (31). This fusion approach assumes that EEG and fTCD evidences are independent with an unequal contribution in correctly inferring user intent.

## 2 Materials and Methods

### 2.1 Data Acquisition

EEG data were collected using a g.tec system with 16 electrodes placed at the positions Fp1, Fp2, F3, F4, Fz, Fc1, Fc2, Cz, P1, P2, C1, C2, Cp3, Cp4, P5, and P6 with the reference electrode being placed at left mastoid. The EEG data were sampled at 256 samples/sec and filtered using the g.tec amplifier’s bandpass (corner frequencies: 2 and 62 Hz) and notch (corner frequencies: 58 and 62 Hz) filters. fTCD data were collected using a SONARA TCD system with two 2 MHz transducers placed at left and right sides of the transtemporal window which is located above the zygomatic arch. fTCD data were recorded at a sampling rate of 44.1 kHz and downsampled by a factor of 5 using low pass filter of 4.4 kHz corner frequency to be 8.82 kHz. A total of 21 healthy participants were included in the study. Each participant completed a single 25-minute session, and all provided a written informed consent prior to the experiment. University of Pittsburgh local Institutional Review Board (IRB) approved all experimental procedures under IRB number of PRO16080475. Data collection took place between April 17 and September 22, 2017. The flickering MR/WG paradigm dataset includes data recorded from 11 healthy subjects (3 females) with age range from 25 to 32 years while MI paradigm dataset includes data recorded from 10 right-handed healthy subjects (6 females) with age range from 23 to 32 years.

### 2.2 Experimental design

The multimodal BCI system employs two visual presentation paradigms to induce simultaneous changes in EEG and fTCD recorded signals. The first paradigm uses motor imagery (MI) tasks while the second one uses flickering mental rotation (MR) and word generation (WG) tasks. Figure 1.A shows the MI paradigm visual presentation (31) with three icons presented on the screen including left and right horizontal arrows representing left arm MI and right arm MI tasks respectively, and a fixation cross representing resting state. When left arm MI task is selected by the vertical red arrow, participants imagine moving left arm. Similarly, right arm movement is imagined if right arm MI task is selected. During the experiment, the vertical arrow points randomly to one of the three icons for 10 s (trial duration) and participants preform the task specified by the vertical arrow. A total of 150 trials were presented to each participant. Figure 2.B shows the flickering MR/WG paradigm visual presentation (29) with three icons presented on the screen. The left icon is a random letter representing the WG task. When selected by the vertical red arrow, participants think of words that begin with the letter displayed on the screen. The right icon shows identical 3D shapes rotated with different angles representing the MR task. When selected by the vertical arrow, participants mentally rotate the shapes to decide if they are identical or mirrored. Finally, the icon in the middle is a fixation cross representing resting state. MR and WG tasks can be distinguished through fTCD due to the differences in blood perfusion they yield on the two sides of the brain (27). Because these tasks do not induce differences in EEG signal, MR/WG tasks were modified to induce differences in EEG signal by being textured with a flickering checkerboard pattern as shown in Figure 1.B. This flickering pattern induces a steady-state visually evoked potential (SSVEPs) in EEG. MR task was modified to flicker at 7 Hz while the WG task was modified to flicker at 17 Hz. Similar to the MI paradigm, the vertical arrow points randomly to one of the three icons for 10 s (trial duration) with a total of 150 trials presented to each participant.

**Figure 1.**
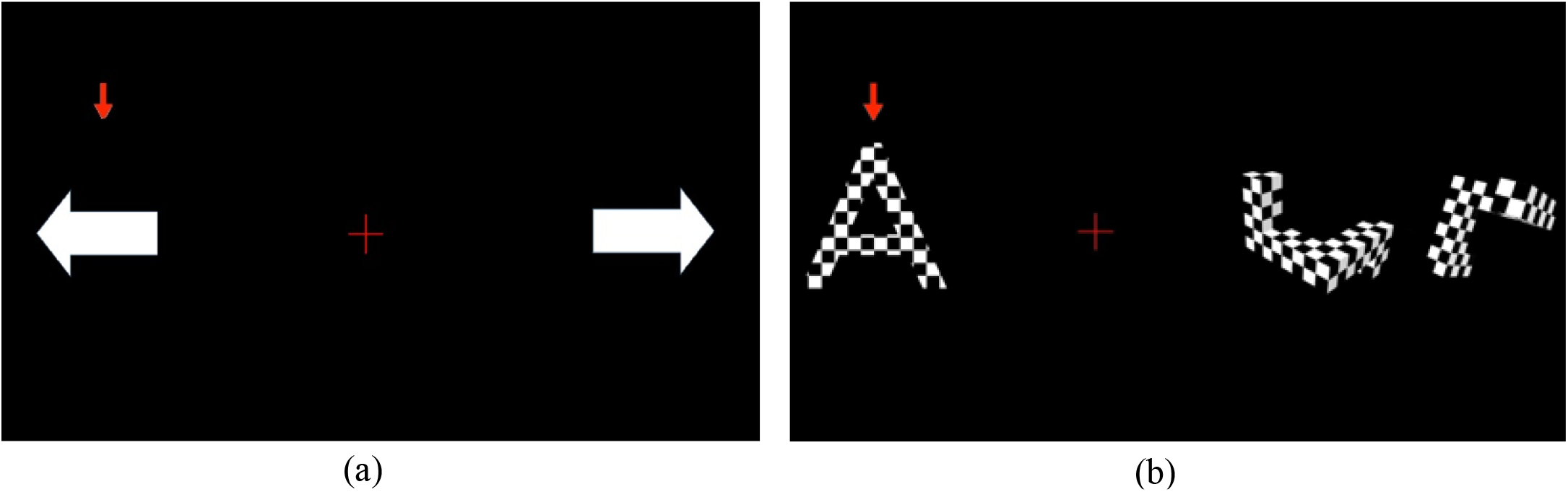
Stimulus presentation for motor imagery EEG-fTCD BCI (a) and flickering MR/WG EEG-fTCD BCI (b).

**Figure 2.** fTCD envelope calculation: (a) raw fTCD signal of a sample trial, (b) spectrogram of raw fTCD signal, and (c) the calculated envelope signal.

### 2.3 EEG Feature Extraction

Common spatial pattern (CSP) is commonly used to extract features in MI EEG-based BCIs (38). CSP computes spatial filters to linearly transform EEG measurements into a space where observations belonging to two classes are more differentiable in terms of variance (39). Recently, filter bank common spatial pattern (FBCSP) was proposed as an extension for CSP (36). To achieve optimal CSP performance, several subject specific parameters have to be specified such as the CSP filters to be used and the operating frequency range to select (40). In a previous study, it was found that CSP features are not significantly differentiable across MI tasks when EEG was not filtered or filtered with an inappropriately selected operating frequency range (41). The FBCSP algorithm overcomes the limitations of the CSP algorithm and eliminates the need to manually select subject-specific frequency range (35,39). In this study, FBCSP was applied to EEG data of both paradigms. While FBCSP is well-known as a feature extractor for MI EEG data, in this work, we extend FBCSP applications and show that it can be a successful feature extraction method when applied to SSVEP MR/WG and MI EEG data. FBCSP applies band-pass filtering on the observations and then extracts CSP features from each frequency band. In particular, the EEG signals from both paradigms were bandpass filtered the signals in the 2-60 Hz range to ensure the SSVEP related changes in the MR/WG paradigm as well the event-related synchronization and desynchronization rhythms in the MI paradigms are fully represented in the EEG signal (42,43). This 2-60 Hz frequency range was divided into 9 non-overlapping frequency bands where each band has a bandwidth of approximately 6.5 Hz. Within each frequency band, CSP finds optimal spatial filters through solving the equation below (44,45):

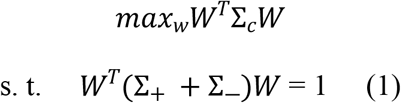

where Σ_*c*_is the average trial covariance matrix for class c ∈ {+, -} and *W*^*T*^Σ_*c*_*W* is the variance in direction W. EEG signal of each trial can be repressed by a matrix *E*^*N*∗*T*^where N is the number of channels and T is the number of samples per channel.

Covariance C of each trial is calculated as:

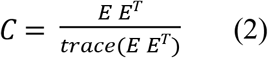

The average covariance per class is given by:

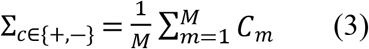

Where M is the number of trials in class c.

Simultaneous diagonalization of Σ_*c*_matrices can find the optimal spatial filters matrix *W* from Equation (1).

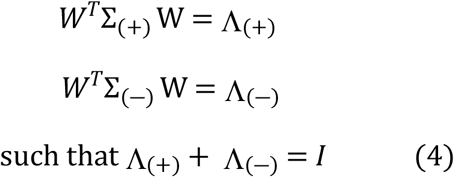

where Λ_c_ is the eigenvalue diagonal matrix. Solution of (4) is equivalent to solution the generalized eigenvalue problem in (5)

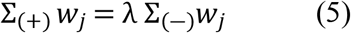

Where *w*_*j*_ is the *j*^*th*^ generalized eigenvector and 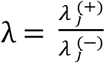

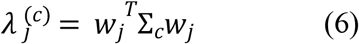

Where 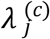 are the diagonal elements of *A* _*c*_. 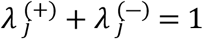 given that Λ_(+)_+ Λ_(−)_ = *I*. Therefore, higher value of 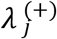 reflects low value of 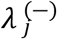 and will result in higher variance in the positive class after filtering using the spatial filter *w*_*j*_ and simultaneously will result in lower variance in the negative class after filtering using the same spatial filter *w*_*j*_.

We solved the binary classification problems of both paradigms at all possible numbers of eigenvectors. In particular, we spatially filtered EEG data using 1, 2, 3, …., and 8 eigenvectors from both ends of W. Therefore, performance of both EEG only and multimodal combination was evaluated using 2*N*_*f*_(2, 4, 6, …., and 16) eigenvectors for each frequency band. CSP features included the log variance of each spatially filtered signal. This yielded 2*N*_*f*_EEG features per band. Features calculated for EEG bands were concatenated to form the EEG features vector which contained 9×2 *N*_*f*_features per trial.

### 2.4 fTCD Feature Extraction

Studies performing fTCD signals analysis commonly extract features from the raw fTCD data (27,37). In this study, we extracted features from the fTCD envelope signal which is derived from the raw fTCD signal captured by the transducers. fTCD envelope signal represents maximal blood flow velocity while raw fTCD signal represents the echoes recorded by the transducers due to many scatterers moving with different velocities (37). To convert the amplitudes of the raw fTCD signal to velocities, the Doppler effect equation below was used.

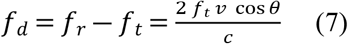

Where *f*_*t*_ is the transmitted frequency, *f*_r_ is the received frequency, *f*_*d*_ is the doppler shift due to the velocity of the scatterers, c is the speed of sound in tissue (1560 cm/s), v is the velocity of the scatterer, and θ is the angle between the ultrasound wave and the flow direction. From this equation, it can be noted that scatterers moving with the highest speed cause the maximal frequency shift. To calculate the envelope signal from raw fTCD data, short-time Fourier transform (STFT) is used to obtain the spectrogram of the raw signal and the maximal frequency which corresponds to the highest blood flow velocity. *f*_*d*_ is plugged into the doppler equation to obtain the corresponding velocity. Figure 2 details the process of calculating the envelope for a sample raw fTCD signal of one trial acquired from the left middle cerebral artery of a single subject.

Time series feature extraction library (TSFEL) was employed to extract features from fTCD envelope signals. The library has been used to extract statistical features for wavelet coefficients as these features have been proved to be successful in fTCD-only BCIs (27). Moreover, we used the library to calculate various sets of features including statistical, temporal features, and spectral domain features. A complete list of features computed by the algorithm is summarized in Figure 3.

**Figure 3.**
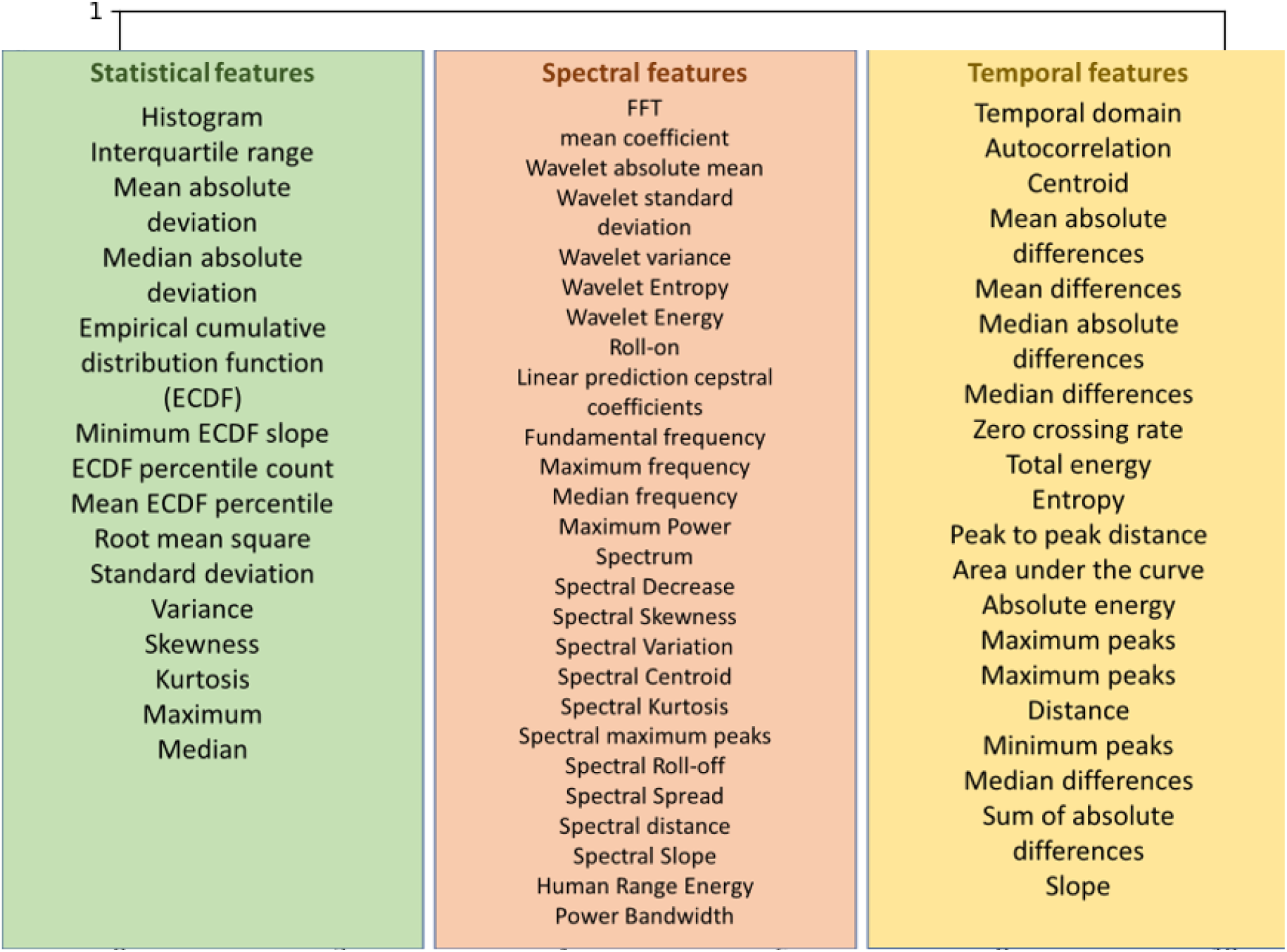
A complete list of features provided by TSFEL.

### 2.5 Feature selection and classification

The Wilcoxon rank-sum test (48) was used to select significant features from fTCD feature vectors of both the MI and MR/WG paradigms at p-value of 0.005. Support vector machine (SVM) classifier with linear kernel (49) was used to evaluate the performance of single-modal BCIs, i.e., EEG only and fTCD only of both paradigms using 10-fold cross validation scheme. fTCD only performance was evaluated using features selected at p-value of 0.005 while EEG only performance was evaluated using 2*N*_*f*_ (2, 4, 6, …., and 16) CSP features per frequency band. fTCD only accuracies as well as best EEG only accuracies were reported in the results section below.

To assess the multimodal BCI performance, SVM was used to project EEG features of each trial into 1-D scalar score (evidence). Similarly, another SVM was used to project fTCD features of each trial into 1-D scalar score (evidence). The fTCD scores were evaluated using fTCD features selected at a p-value of 0.05 while the EEG scores were evaluated using 2*N*_*f*_ (2, 4, 6,, and 16) CSP features per frequency band. Bayesian fusion was used to infer user intent based on EEG and fTCD scores (evidences). Best multimodal accuracies were reported in the results section below.

### 2.6 Bayesian Probabilistic Fusion

To generate a joint decision of a test trial based on information from both modalities, we performed a probabilistic Bayesian fusion of EEG and fTCD evidences obtained from the training trials under the assumption that these evidences come from independent distributions, and may have equal or unequal weight in user intent inference (31).

#### 2.6.1 Weighted Independent Probabilistic Fusion

The 1-D SVM scores generated from each modality, also called evidences, are split into training and testing sets using 10-fold cross validation. The goal is to infer the user intent *X*_*k*_ of a test trial given a set of paired evidences *Y* = {*y*1,*y*_2,_…*y*_*N*―10_} obtained from the training data where N is the number of trials, and each element of this set *y*_*n*_ = {*e*_*n*_,*f*_*n*_} represents the EEG (*e*_*n*)_ and fTCD (*f*_*n*_) evidences of one trial. The user intent *X*_*k*_ of a test trial *y*_*k*_ = {*e*_*k*_,*f*_*k*_} is determined through joint state estimation using EEG and fTCD evidences as follows:

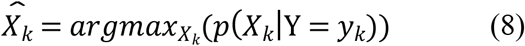

where *p*(*X*_*k*_|Y) is the state posterior distribution conditioned on Y. Using Bayes rule, (8) can be formulated as:

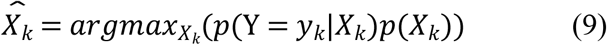

where *p*(Y|*X*_*k*_) is the state conditional distribution of Y and *p*(*X*_*k*_) is the prior distribution of *X*_*k*_. Since the trials are randomized, the prior distribution is assumed to be uniform. Consequently, (9) can be written as:

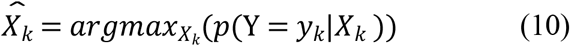

The distribution *p*(Y|*X*_*k*_)) can be computed Using EEG and fTCD evidences of the training trials.

Assuming that the EEG and fTCD evidences conditioned on *X*_*k*_ are independent, (10) can be written as follows:

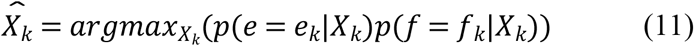

The distributions *p*(*e*|*X*_*k*_) and *p*(*f*|*X*_*k*_) represent the EEG and fTCD evidences distributions conditioned on *X*_*k*_, and are computed in each fold from the N-10 training scores using kernel density estimation with a gaussian kernel and Scott’s rule of thumb as the bandwidth selector. The probabilities *p*(*e* = *e*_*k*_|*X*_*k*_) and *p*(*f* = *f*_*k*_|*X*_*k*_) for each test trial are computed from the distributions and plugged in equation (11), and the decision 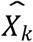 that maximizes the likelihood is selected. Equation 15 was modified to allow for the possibility that EEG and fTCD evidences does not have equal contribution in decision making, yielding:

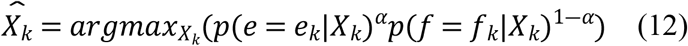

where *α* is a weighting factor determined via a grid search from 0 to 1 with a step of 0.01.

## 3 Results

In this section, we evaluate the performance of the MI and MR/WG EEG-fTCD systems when employing the proposed analysis pipeline which includes FBCSP for EEG feature extraction and time series features for fTCD envelope feature extraction with weighted Bayesian fusion for multimodal decision making. The following experimental results provide accuracy comparisons of the MI and MR/WG multimodal BCIs against the highest performance achieved using EEG-only and fTCD-only BCIs.

### 3.1 Motor Imagery Paradigm

Table 1 shows maximum accuracy achieved for each subject in right MI versus baseline, left MI versus baseline, and right MI versus left MI selection problems respectively using weighted probabilistic fusion and the corresponding accuracies using EEG only and fTCD only. In order to assess the significance of the multimodal model, one-sided paired Wilcoxon signed rank test was used to statistically compare the accuracies of the multimodal system with the accuracies obtained using EEG only. Table 1 demonstrates that the average accuracies for right MI versus baseline selection problem are 90.73% for EEG only, 51.98% for fTCD only, and 94.53% for the weighted probabilistic fusion. Accuracies obtained using the weighted probabilistic fusion model are statistically significant compared to those obtained using EEG only with a p-value of 0.042 (Table 2). Left MI versus baseline performance measures in Table 1 show average accuracies of 91.03% and 52.58% for EEG only and fTCD only respectively while the weighted probabilistic fusion achieved 94.9%. The weighted probabilistic fusion model resulted in a statistically significant increase in accuracy with a p -value of 0.0098 compared to EEG only as shown in Table 2. The third task, right MI versus left MI, shows an average accuracy of 96.29 % for the weighted probabilistic fusion which outperforms the accuracies of 90.48% and 51.43% obtained using EEG only and fTCD only respectively. In comparison with EEG only, the weighted probabilistic fusion model shows significance with a p-value of 0.002 as shown in Table 2.

**Table 1.** Maximum accuracy achieved for each subject using weighted fusion and the corresponding accuracies obtained using EEG only and fTCD only for MI paradigm.

|  | Sub_ID | 1 | 2 | 3 | 4 | 5 | 6 | 7 | 8 | 9 | 10 | Mean |
| --- | --- | --- | --- | --- | --- | --- | --- | --- | --- | --- | --- | --- |
| Baseline vs Left | EEG | 96.91 | 95.88 | 80.41 | 94.85 | 97.94 | 91.75 | 83.51 | 88.66 | 91.75 | 88.66 | 91.03 |
|  | ftCD | 69.71 | 49.48 | 42.27 | 58.76 | 57.73 | 57.73 | 48.45 | 45.36 | 47.42 | 49.48 | 52.58 |
|  | Fusion | 97.92 | 96.88 | 95.83 | 94.79 | 96.88 | 95.83 | 88.54 | 95.83 | 95.83 | 90.62 | 94.9 |
| Baseline vs Right | EEG | 98.96 | 90.62 | 91.67 | 81.25 | 95.83 | 97.92 | 83.33 | 89.58 | 94.79 | 83.33 | 90.73 |
|  | ftCD | 48.96 | 46.88 | 48.96 | 66.67 | 46.88 | 60.42 | 51.04 | 48.96 | 38.54 | 62.5 | 51.98 |
|  | Fusion | 97.89 | 94.74 | 91.58 | 92.63 | 96.84 | 96.84 | 89.47 | 97.89 | 95.79 | 91.58 | 94.53 |
| Right vs Left | EEG | 99.05 | 77.14 | 83.81 | 96.19 | 94.29 | 92.38 | 83.81 | 85.71 | 98.1 | 94.29 | 90.48 |
|  | ftCD | 51.43 | 69.52 | 49.52 | 48.57 | 60.95 | 53.33 | 47.62 | 41.9 | 48.57 | 42.86 | 51.43 |
|  | Fusion | 99.05 | 85.71 | 89.52 | 98.1 | 100 | 99.05 | 94.29 | 97.14 | 100 | 100 | 96.29 |

**Table 2.**
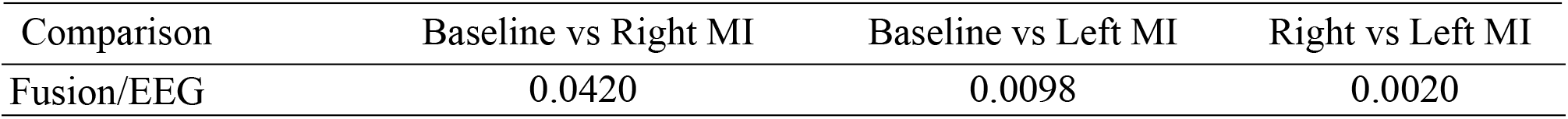
P-values showing accuracy significance of weighted probabilistic fusion compared to EEG only for the MI paradigm.

The optimal alpha values that give the highest accuracy for each subject are reported in Figure 4.A and the average of these values per task is reported in Figure 4.B. These alpha values represent the contribution of each of the two modalities in decision making in the three classification problems. More specifically, alpha is the weighting factor for EEG modality and (1-alpha) is the fTCD weighting factor, so higher alpha values reflect higher EEG contribution in discriminating the tasks at the given problem. To test wither the contribution of EEG and fTCD modalities in decision making is task-dependent, one-sided Wilcoxon rank-sum test with a p-value of 0.05 was applied to the optimal alpha values of each subject within each task to check if their distribution have a median lower or higher than 0.5 where 0.5 represents equal EEG and fTCD contributions. No significance was observed for the three classification problems.

**Figure 4.**
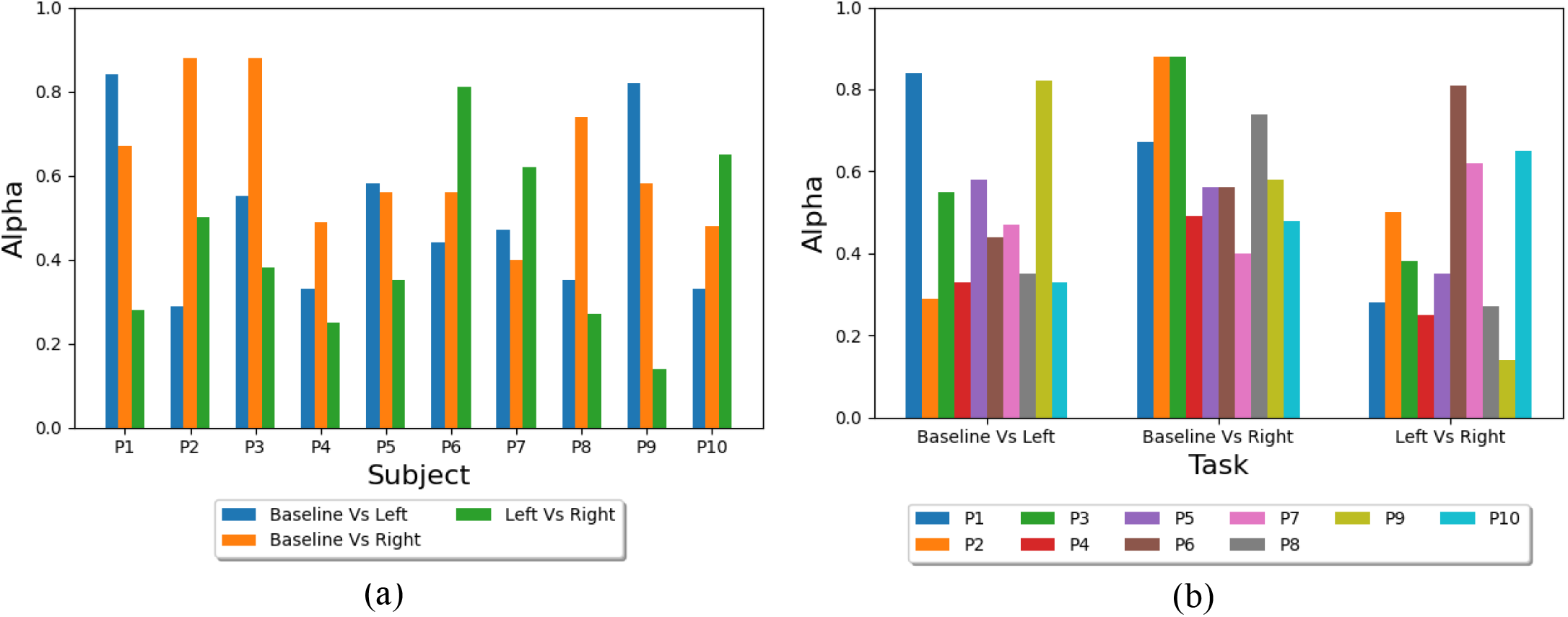
Optimal alpha values: (a) The values of alpha yielding the highest accuracy for each subject in Baseline Vs Left, Baseline Vs Right, and Left Vs Right tasks, and (b) Average values of alpha yielding the highest accuracy for each task (Baseline Vs Left, Baseline Vs Right, and Left Vs Right).

### 3.2 Flickering MR/WG Paradigm

The maximum accuracy achieved per subject using weighted probabilistic fusion and the corresponding EEG only accuracy and fTCD only accuracy are reported in Table 3 for MR versus baseline, WG versus baseline, and MR versus WG problems. Table 4 shows the calculated p-values using one-sided Wilcoxon signed rank test to compare the significance of the weighted probabilistic fusion with EEG only in terms of accuracy. MR versus baseline problem achieved average accuracies of 95.27% for the weighted probabilistic fusion model which is higher than 92.99% for EEG only and 50% for fTCD only (Table 3). Table 4 shows a p-value of 0.0156 describing the statistical significance of the increase in accuracy of the weighted probabilistic fusion in comparison with EEG only. Table 3 also shows the performance measures for WG versus baseline problem. In particular, weighted probabilistic fusion achieved average accuracy of 85.93% compared to 80.98% for EEG only and 52.3 % for fTCD only. Table 4 proves the significance of the weighted probabilistic fusion in terms of accuracy compared to EEG only with a p-value of 0.0049. As for MR versus WG problem, we obtained the highest average accuracy compared to MR/WG versus baseline problems as seen in Table 3. In particular, average accuracy of 96.97% for the weighted probabilistic fusion was obtained which outperformed the average accuracy of 95.41% and 50.82% obtained using EEG only and fTCD only respectively. Table 4 shows a p-value of 0.0469 for the weighted fusion which indicates that it is statistically significant compared to EEG only.

**Table 3.** Maximum accuracy achieved for each subject using weighted fusion and the corresponding accuracies obtained using EEG only and fTCD only for flickering MR/WG paradigm.

|  | Sub_ID | 1 | 2 | 3 | 4 | 5 | 6 | 7 | 8 | 9 | 10 | 11 | Mean |
| --- | --- | --- | --- | --- | --- | --- | --- | --- | --- | --- | --- | --- | --- |
| Baseline vs MR | EEG | 92.71 | 96.88 | 88.54 | 88.54 | 91.67 | 89.58 | 91.67 | 97.92 | 98.96 | 95.83 | 90.62 | 92.99 |
|  | ftCD | 41.67 | 44.79 | 55.21 | 59.38 | 63.54 | 39.58 | 48.96 | 44.79 | 59.38 | 41.67 | 51.04 | 50.0 |
|  | Fusion | 93.75 | 96.88 | 89.58 | 91.67 | 96.88 | 95.83 | 96.88 | 96.88 | 98.96 | 95.83 | 94.79 | 95.27 |
| Baseline vs WG | EEG | 89.69 | 85.57 | 83.51 | 76.29 | 73.2 | 83.51 | 81.44 | 91.75 | 94.85 | 59.79 | 71.13 | 80.98 |
|  | ftCD | 58.76 | 55.67 | 63.92 | 50.52 | 71.13 | 50.52 | 42.27 | 39.18 | 57.73 | 42.27 | 43.3 | 52.3 |
|  | Fusion | 91.58 | 92.63 | 89.47 | 72.63 | 85.26 | 86.32 | 87.37 | 93.68 | 95.79 | 67.37 | 83.16 | 85.93 |
| MR vs WG | EEG | 96.19 | 99.05 | 91.43 | 87.62 | 97.14 | 99.05 | 87.62 | 98.1 | 100 | 96.19 | 97.14 | 95.41 |
|  | ftCD | 49.52 | 49.52 | 45.71 | 49.52 | 44.76 | 57.14 | 66.67 | 44.76 | 56.19 | 41.9 | 53.33 | 50.82 |
|  | Fusion | 97.14 | 99.05 | 98.1 | 86.67 | 97.14 | 100 | 95.24 | 98.1 | 99.05 | 98.1 | 98.1 | 96.97 |

**Table 4.** P-values showing accuracy significance of weighted fusion compared to EEG only for flickering MR/WG paradigm.

| Comparisons | Baseline vs MR | Baseline vs WG | MR vs WG |
| --- | --- | --- | --- |
| Fusion/EEG | 0.0156 | 0.0049 | 0.0469 |

The optimal alpha values for each subject are reported in Figure 5.A and the average of these values per task is reported in Figure 5.B. Statical significance testing similar to the one performed in section 3.1 was applied to the alpha values of each task. Baseline versus MR and baseline versus WG tasks showed no significance while the alpha values of MR versus WG task was significantly lower than 0.5 with a p-value of 0.03.

**Figure 5.**
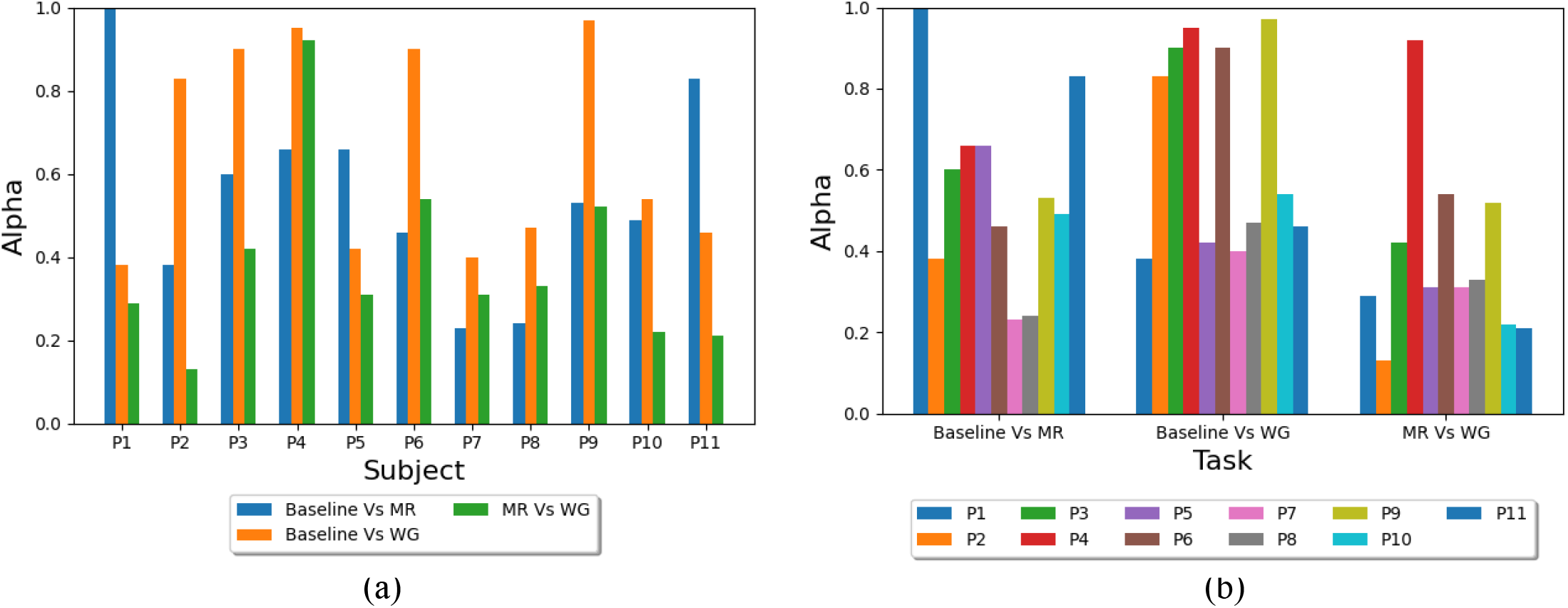
Optimal alpha values: (a) The values of alpha yielding the highest accuracy for each subject in Baseline Vs MR, Baseline Vs WG, and MR Vs WG tasks, and (b) Average values of alpha yielding the highest accuracy for each task (Baseline Vs MR, Baseline Vs WG, and MR Vs WG).

## 4 Discussion

To enhance the performance of MI and MR/WG paradigms, we employed FBCSP to analyze EEG data of both paradigms and extracted time series features from the envelope of fTCD signals of both paradigms. Probabilistic Bayesian fusion was used to infer user intent based on EEG and fTCD input. Probabilistic fusion obtained statistically significant higher accuracies than EEG only by 3.87%, 3.80%, and 5.81% on average for baseline versus left MI, baseline versus right MI, and right MI versus left MI respectively (Table 1). Interestingly, FBCSP which is known as a successful feature extraction method for MI-based BCIs, yielded high accuracy when used to analyze EEG data of the MR/WG SSVEP paradigm. In fact, performance measures obtained with FBCSP outperform the feature extraction methods we used in previous studies for MR/WG paradigm (29,32). Average accuracies of 95.27%, 85.93%, and 96.97% were obtained for MR versus baseline, WG versus baseline, and MR versus WG respectively using Bayesian fusion with average increases of 2.28%, 4.95%, and 1.56% compared to EEG only (Table 2). It can be noted that the average accuracy for the WG versus baseline problem is much lower than the average accuracies of MR versus baseline and WG versus baseline. The reason for such accuracy drop is unknown based on the available data, but requires further investigations. Although fTCD only performance accuracy was low for both paradigms, it was able to improve the multimodal performance when combined with EEG. Moreover, using the envelope fTCD signal instead of the raw signal has significantly sped up computations, especially in feature extraction, as the envelope signal has much lower dimensionality. This increase in computational efficiency made it possible to calculate several sets of features that were impossible to be calculated on the raw fTCD signal due to their computational complexity. Contribution of each modality in user intent inference was investigated per task and it was found that EEG has significantly lower contribution than fTCD in WG versus MR task. Other tasks did not show higher contribution for any of the modalities. Larger sample size is necessary to further investigate EEG and fTCD contributions across tasks.

Table 5 shows a comparison between EEG-fTCD performance obtained using the proposed analysis approach and the performance we obtained in previous studies (31,32). For task versus baseline problems in both paradigms, similar or higher accuracies were obtained with the current analysis pipeline compared to those we obtained previously. In particular, average accuracies of 95.27% and 85.93% were obtained for MR versus baseline and WG versus baseline problems respectively with the proposed analysis approach compared to 86.27% and 85.29% obtained previously. In addition, accuracies of 94.53% and 94.9% were achieved for left MI versus baseline and right MI versus baseline respectively compared to 93.71% and 93.85% obtained in previous studies. However, the accuracy of the proposed approach dropped by 1.2% and 3.7% for MR versus WG and left MI versus right MI respectively compared to our previous results. Despite the drop in task versus task accuracy, we believe the current analysis approach is more successful than the approaches we introduced earlier especially for MR/WG paradigm as it led to a significant 9% increase in performance accuracy for the MR versus baseline problem.

**Table 5.** Comparison between the EEG-fTCD BCI performance with proposed analysis approach and the approaches introduced previously.

| Method | Modality | Activity | Accuracy | Task-Baseline/rest length (s) |
| --- | --- | --- | --- | --- |
| (31) Khalaf et al.,2019 (Left/baseline) | EEG+fTCD | Motor Imagery | 93.71% | 10 – NA |
| (31)Khalaf et al.,2019 (Right/baseline) | EEG+fTCD | Motor Imagery | 93.85% | 10 – NA |
| (31)Khalaf et al.,2019 (Left/Right) | EEG+fTCD | Motor Imagery | 100.00% | 10 – NA |
| (32)Khalaf et al.,2019 (MR/baseline) | EEG+fTCD | Flickering MR/WG | 86.27% | 10 – NA |
| (32) Khalaf et al., 2019 (WG/baseline) | EEG+ftCD | Flickering<br>MR/WG | 85.29% | 10 – NA |
| (32)Khalaf et al., 2019 (MR/WG) | EEG+ftCD | Flickering<br>MR/WG | 98.18% | 10 – NA |
| Proposed Method (Left/baseline) | EEG+ftCD | Motor<br>Imagery | 94.53% | 10 – NA |
| Proposed Method (Right/baseline) | EEG+ftCD | Motor<br>Imagery | 94.9% | 10 – NA |
| Proposed Method (Left/Right) | EEG+ftCD | Motor<br>Imagery | 96.29% | 10 – NA |
| Proposed Method (MR/baseline) | EEG+ftCD | Flickering<br>MR/WG | 95.27% | 10 – NA |
| Proposed Method (WG/baseline) | EEG+ftCD | Flickering<br>MR/WG | 85.93% | 10 – NA |
| Proposed Method (MR/WG) | EEG+ftCD | Flickering<br>MR/WG | 96.97% | 10 – NA |

Moreover, when comparing the performance of our system with the performance of EEG-fNIRS BCI as shown in Table 6, it can be noted that the performance of the three binary selection problems within the MI paradigm as well as two out of the three selection problems within the MR/WG paradigm outperforms all the multimodal EEG-fNIRS BCIs in terms of accuracy. In addition, our system has the shortest trial length compared to all the multimodal BCIs in comparison expect for the BCI by Buccino et al. (50), however, that BCI requires a rest period of 6 seconds in addition to 6 seconds trial length while our system does not require any rest periods between trials. In fact, the rest task in our system (when the user focuses on the fixation cross) is randomly selected by the vertical arrow in the visual paradigm and is considered a separate task resembling the situation when the subject does not want to issue any commands while in the other BCIs in comparison, each trial is followed by a rest period to stabilize the hemodynamic response before the next trial. Moreover, the study that achieved the highest EEG-fNIRS accuracy of 94.2% used a motor execution task (50) while our system does not require any muscular input from the users as it is intended to be used for patients with severe motor disabilities. The 85.93% accuracy obtained by WG versus baseline problem is in fact worse that the accuracies obtained by several EEG-fNRIS systems (50, 51,52,55,56). However, the other systems are either much slower (trial duration + rest period) or require muscular input.

**Table 6.**
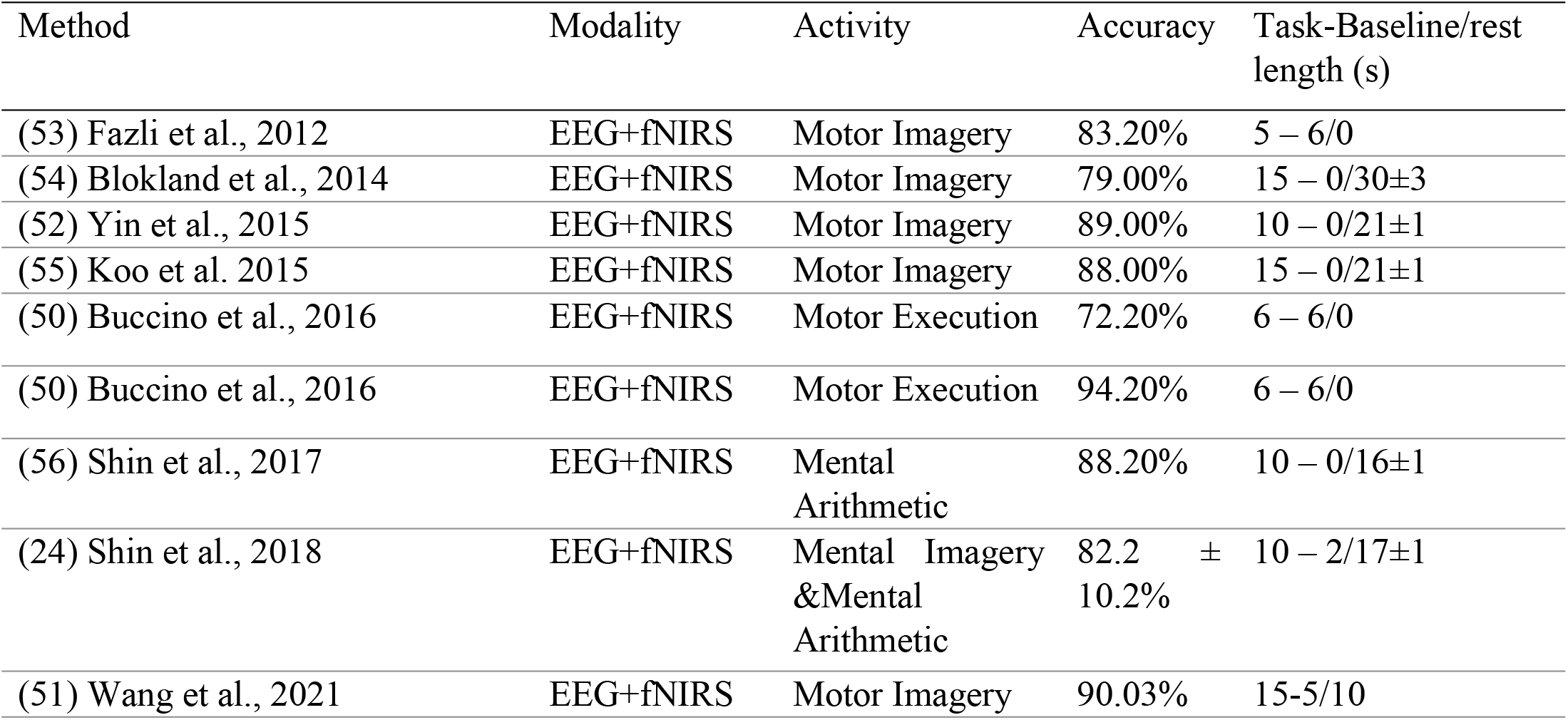
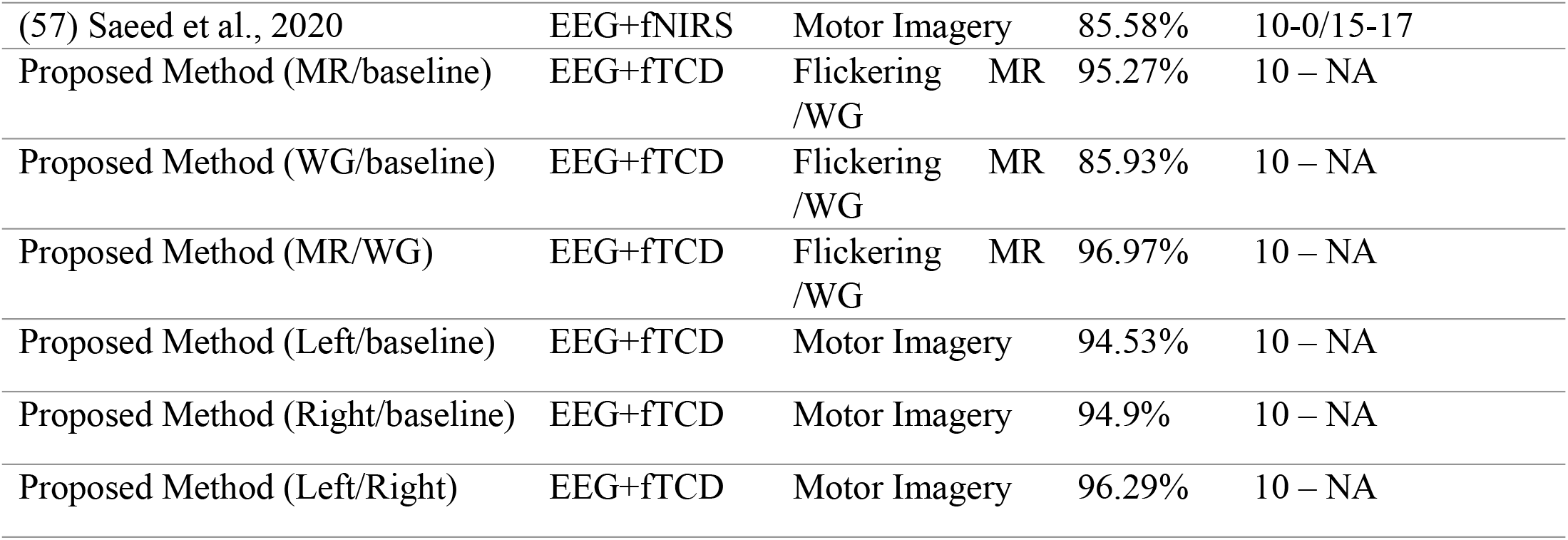
Comparison between MI and flickering WG/MR EEG-fTCD multimodal BCIs with the proposed analysis approach and the state-of-the-art multimodal EEG-fNIRS in literature.

One limitation of the proposed analysis pipeline is the low performance accuracy of fTCD only. Future directions include investigating more features of fTCD envelope and fTCD raw signals. This is expected to boost the fTCD accuracy and thus the multimodal performance. Another limitation is the lack of explanation of the accuracy drop in WG versus baseline problem compared to the other selection problems in the MR/WG paradigm. Larger sample size is needed to investigate the differences in contribution of each modality in user intent inference across tasks and paradigms. Other future directions include calibration time reduction via invariant representation learning deep networks. Discovering and exploiting shared invariant neural activity in EEG-fTCD joint space is of significant interest for enabling generalizability of decoding models across subjects and recording sessions.

## 5 Conclusion

In this paper, we employed FBCSP and time series features to analyze EEG signals and envelope of fTCD signals in multimodal EEG-fTCD BCIs. Moreover, instead of concatenating EEG and fTCD feature vectors of each trial, we employed a probabilistic fusion approach of EEG and fTCD evidences assuming that these evidences are independent, but with unequal contribution in inferring user intent. Binary classification problems were investigated for both MI and flickering MR/WG paradigms. For MR/WG paradigm, the multimodal system achieved average accuracies of 95.27 %, 85.98% and 96.97% for baseline versus MR, baseline versus WG, and MR versus WG respectively. As for the MI paradigm, average accuracies of 94.53%, 94.9% and 96.29% were obtained for baseline versus left, baseline versus right, and left versus right, respectively. The multimodal EEG-fTCD BCI with the proposed analysis pipeline outperformed all EEG-fNRIS BCIs in comparison.

## Author Contributions

A.E., A.E., A.Z., M.E., and S.M. contributed to formal analysis, methodology, data curation, software, validation, visualization, and writing – original draft. A.A contributed to methodology, software, and writing – review and editing. A.E. contributed to methodology and writing – review and editing, M.A. contributed to investigation, resources, and writing – review and editing. A.K. contributed to conceptualization, investigation, methodology, project administration, data curation, supervision, and writing – review and editing.

## References

1. Nicolas-Alonso LF, Gomez-Gil J. Brain computer interfaces, a review. Sensors (Basel). 2012;12(2):1211–79.

2. Khan MU, Hasan MAH. Hybrid EEG-fNIRS BCI Fusion Using Multi-Resolution Singular Value Decomposition (MSVD). Front Hum Neurosci. 2020 Dec 8;14.

3. Lazarou I, Nikolopoulos S, Petrantonakis PC, Kompatsiaris I, Tsolaki M. EEG-Based Brain– Computer Interfaces for Communication and Rehabilitation of People with Motor Impairment: A Novel Approach of the 21st Century. Front Hum Neurosci. 2018 Jan 31;12.

4. Duan X, Xie S, Xie X, Meng Y, Xu Z. Quadcopter Flight Control Using a Non-invasive Multi-Modal Brain Computer Interface. Front Neurorobot. 2019 May 31;13.

5. Yanagisawa T, Fukuma R, Seymour B, Hosomi K, Kishima H, Shimizu T, et al. Using a BCI Prosthetic Hand to Control Phantom Limb Pain. In 2019. p. 43–52.

6. Khan RA, Naseer N, Qureshi NK, Noori FM, Nazeer H, Khan MU. fNIRS-based Neurorobotic Interface for gait rehabilitation. J Neuroeng Rehabil. 2018 Dec 5;15(1):7.

7. Coogan CG, He B. Brain-Computer Interface Control in a Virtual Reality Environment and Applications for the Internet of Things. IEEE Access. 2018;6:10840–9.

8. Ahn M, Lee M, Choi J, Jun S. A Review of Brain-Computer Interface Games and an Opinion Survey from Researchers, Developers and Users. Sensors. 2014 Aug 11;14(8):14601–33.

9. Waldert S. Invasive vs. Non-Invasive Neuronal Signals for Brain-Machine Interfaces: Will One Prevail? Front Neurosci. 2016 Jun 27;10.

10. Kwon J, Shin J, Im CH. Toward a compact hybrid brain-computer interface (BCI): Performance evaluation of multi-class hybrid EEG-fNIRS BCIs with limited number of channels. PLoS One. 2020 Mar 18;15(3):e0230491.

11. Lotte F, Congedo M, Lécuyer A, Lamarche F, Arnaldi B. A review of classification algorithms for EEG-based brain–computer interfaces. J Neural Eng. 2007 Jun 1;4(2):R1–13.

12. Min BK, Marzelli MJ, Yoo SS. Neuroimaging-based approaches in the brain–computer interface. Trends Biotechnol. 2010 Nov;28(11):552–60.

13. Naseer N, Hong KS. fNIRS-based brain-computer interfaces: a review. Front Hum Neurosci. 2015 Jan 28;9.

14. Sokunbi MO, Gradin VB, Waiter GD, Cameron GG, Ahearn TS, Murray AD, et al. Nonlinear Complexity Analysis of Brain fMRI Signals in Schizophrenia. PLoS One. 2014 May 13;9(5):e95146.

15. Lal SKL, Craig A, Boord P, Kirkup L, Nguyen H. Development of an algorithm for an EEG-based driver fatigue countermeasure. J Safety Res. 2003 Aug;34(3):321–8.

16. Aydemir O, Kayikcioglu T. Decision tree structure based classification of EEG signals recorded during two dimensional cursor movement imagery. J Neurosci Methods. 2014 May;229:68–75.

17. Hasan MAH, Khan MU, Mishra D. A Computationally Efficient Method for Hybrid EEG-fNIRS BCI Based on the Pearson Correlation. Biomed Res Int. 2020 Aug 19;2020:1–13.

18. Brandl S, Hohne J, Muller KR, Samek W. Bringing BCI into everyday life: Motor imagery in a pseudo realistic environment. In: 2015 7th International IEEE/EMBS Conference on Neural Engineering (NER). IEEE; 2015. p. 224–7.

19. Hong KS, Khan MJ. Hybrid Brain–Computer Interface Techniques for Improved Classification Accuracy and Increased Number of Commands: A Review. Front Neurorobot. 2017 Jul 24;11.

20. Mano M, Lécuyer A, Bannier E, Perronnet L, Noorzadeh S, Barillot C. How to Build a Hybrid Neurofeedback Platform Combining EEG and fMRI. Front Neurosci. 2017 Mar 21;11.

21. Allison BZ, Wolpaw EW, Wolpaw JR. Brain–computer interface systems: progress and prospects. Expert Rev Med Devices. 2007 Jul 9;4(4):463–74.

22. Buccino AP, Keles HO, Omurtag A. Hybrid EEG-fNIRS Asynchronous Brain-Computer Interface for Multiple Motor Tasks. PLoS One. 2016 Jan 5;11(1):e0146610.

23. Khan MJ, Hong KS. Hybrid EEG–fNIRS-Based Eight-Command Decoding for BCI: Application to Quadcopter Control. Front Neurorobot. 2017 Feb 17;11.

24. Shin J, Kwon J, Im CH. A Ternary Hybrid EEG-NIRS Brain-Computer Interface for the Classification of Brain Activation Patterns during Mental Arithmetic, Motor Imagery, and Idle State. Front Neuroinform. 2018 Feb 23;12.

25. Zephaniah P v., Kim JG. Recent functional near infrared spectroscopy based brain computer interface systems: Developments, applications and challenges. Biomed Eng Lett. 2014 Sep 18;4(3):223–30.

26. Naseer N, Hong KS. fNIRS-based brain-computer interfaces: A review. Front Hum Neurosci. 2015 Jan 28;9(JAN):3.

27. Khalaf A, Sybeldon M, Sejdic E, Akcakaya M. A brain-computer interface based on functional transcranial doppler ultrasound using wavelet transform and support vector machines. J Neurosci Methods. 2018 Jan;293:174–82.

28. Alexandrov A v., Sloan MA, Wong LKS, Douville C, Razumovsky AY, Koroshetz WJ, et al. Practice standards for transcranial Doppler ultrasound: part I--test performance. J Neuroimaging [Internet]. 2007 Jan [cited 2022 Aug 24];17(1):11–8. Available from: https://pubmed.ncbi.nlm.nih.gov/17238867/

29. Khalaf A, Sejdic E, Akcakaya M. Towards optimal visual presentation design for hybrid EEG—fTCD brain–computer interfaces. J Neural Eng. 2018 Oct 1;15(5):056019.

30. Khalaf A, Sejdic E, Akcakaya M. A novel motor imagery hybrid brain computer interface using EEG and functional transcranial Doppler ultrasound. J Neurosci Methods. 2019 Feb;313:44–53.

31. Khalaf A, Sejdic E, Akcakaya M. Common spatial pattern and wavelet decomposition for motor imagery EEG-fTCD brain-computer interface. J Neurosci Methods. 2019 May 15;320:98–106.

32. Khalaf A, Sejdic E, Akcakaya M. EEG-fTCD hybrid brain–computer interface using template matching and wavelet decomposition. J Neural Eng. 2019 Jun 1;16(3):036014.

33. Khalaf A, Sejdic E, Murat A. Three-Class EEG-fTCD Brain-Computer Interfaces. 2020 Aug 29 [cited 2022 Aug 24]; Available from: /articles/preprint/Three-Class_EEG-fTCD_Brain-Computer_Interfaces/12867260/1

34. Aggarwal S, Chugh N. Signal processing techniques for motor imagery brain computer interface: A review. Array. 2019 Jan;1–2:100003.

35. Ang KK, Chin ZY, Wang C, Guan C, Zhang H. Filter Bank Common Spatial Pattern Algorithm on BCI Competition IV Datasets 2a and 2b. Front Neurosci. 2012;6.

36. Kai Keng Ang, Zhang Yang Chin, Haihong Zhang, Cuntai Guan. Filter Bank Common Spatial Pattern (FBCSP) in Brain-Computer Interface. In: 2008 IEEE International Joint Conference on Neural Networks (IEEE World Congress on Computational Intelligence). IEEE; 2008. p. 2390–7.

37. Sejdić E, Kalika D, Czarnek N. An Analysis of Resting-State Functional Transcranial Doppler Recordings from Middle Cerebral Arteries. PLoS One. 2013 Feb 6;8(2):e55405.

38. Devlaminck D, Wyns B, Grosse-Wentrup M, Otte G, Santens P. Multisubject Learning for Common Spatial Patterns in Motor-Imagery BCI. Comput Intell Neurosci. 2011;2011:1–9.

39. Blankertz B, Tomioka R, Lemm S, Kawanabe M, Muller K robert. Optimizing Spatial filters for Robust EEG Single-Trial Analysis. IEEE Signal Process Mag. 2008;25(1):41–56.

40. Blankertz B, Tomioka R, Lemm S, Kawanabe M, Muller K robert. Optimizing Spatial filters for Robust EEG Single-Trial Analysis. IEEE Signal Process Mag. 2008;25(1):41–56.

41. Novi Q, Guan C, Dat TH, Xue P. Sub-band Common Spatial Pattern (SBCSP) for Brain-Computer Interface. In: 2007 3rd International IEEE/EMBS Conference on Neural Engineering. IEEE; 2007. p. 204–7.

42. Vansteensel MJ, Jarosiewicz B. Brain-computer interfaces for communication. In 2020. p. 67–85.

43. Ha KW, Jeong JW. Motor Imagery EEG Classification Using Capsule Networks. Sensors. 2019 Jun 27;19(13):2854.

44. Ai Q, Chen A, Chen K, Liu Q, Zhou T, Xin S, et al. Feature extraction of four-class motor imagery EEG signals based on functional brain network. J Neural Eng. 2019 Apr 1;16(2):026032.

45. Tan P, Wang X, Wang Y. Dimensionality reduction in evolutionary algorithms-based feature selection for motor imagery brain-computer interface. Swarm Evol Comput. 2020 Feb;52:100597.

46. Grosse-Wentrup M, Liefhold C, Gramann K, Buss M. Beamforming in Noninvasive Brain– Computer Interfaces. IEEE Trans Biomed Eng. 2009 Apr;56(4):1209–19.

47. Haiping Lu, How-Lung Eng, Cuntai Guan, Plataniotis KN, Venetsanopoulos AN. Regularized Common Spatial Pattern With Aggregation for EEG Classification in Small-Sample Setting. IEEE Trans Biomed Eng. 2010 Dec;57(12):2936–46.

48. Rey D, Neuhäuser M. Wilcoxon-Signed-Rank Test. In: International Encyclopedia of Statistical Science. Berlin, Heidelberg: Sprimger Berlin Heidelberg; 2011. p. 1658–9.

49. Chih-Wei Hsu, Chih-Jen Lin. A comparison of methods for multiclass support vector machines. IEEE Trans Neural Netw. 2002 Mar;13(2):415–25.

50. Buccino AP, Keles HO, Omurtag A. Hybrid EEG-fNIRS Asynchronous Brain-Computer Interface for Multiple Motor Tasks. PLoS One. 2016 Jan 5;11(1):e0146610.

51. Wang P, He J, Lan W, Yang H, Leng Y, Wang R, et al. A Hybrid EEG-fNIRS Brain-Computer Interface Based on Dynamic Functional Connectivity and Long Short-Term Memory. In: 2021 IEEE 5th Advanced Information Technology, Electronic and Automation Control Conference (IAEAC). IEEE; 2021. p. 2214–9.

52. Yin X, Xu B, Jiang C, Fu Y, Wang Z, Li H, et al. A hybrid BCI based on EEG and fNIRS signals improves the performance of decoding motor imagery of both force and speed of hand clenching. J Neural Eng. 2015 Jun 1;12(3):036004.

53. Fazli S, Mehnert J, Steinbrink J, Curio G, Villringer A, Müller KR, et al. Enhanced performance by a hybrid NIRS–EEG brain computer interface. Neuroimage. 2012 Jan;59(1):519–29.

54. Blokland Y, Spyrou L, Thijssen D, Eijsvogels T, Colier W, Floor-Westerdijk M, et al. Combined EEG-fNIRS Decoding of Motor Attempt and Imagery for Brain Switch Control: An Offline Study in Patients With Tetraplegia. IEEE Transactions on Neural Systems and Rehabilitation Engineering. 2014 Mar;22(2):222–9.

55. Koo B, Lee HG, Nam Y, Kang H, Koh CS, Shin HC, et al. A hybrid NIRS-EEG system for self-paced brain computer interface with online motor imagery. J Neurosci Methods. 2015 Apr;244:26–32.

56. Shin J, Müller KR, Schmitz CH, Kim DW, Hwang HJ. Evaluation of a Compact Hybrid Brain-Computer Interface System. Biomed Res Int. 2017;2017:1–11.

57. Saeed A, Naseer N, Jabbar H. Improving classification performance of hybrid EEG-fNIRS BCI system by channel optimization. In: Proceedings of the 13th ACM International Conference on PErvasive Technologies Related to Assistive Environments. New York, NY, USA: ACM; 2020. p. 1–4.

